# Central Vision Loss Alters Internal Emotion Representations: Evidence from a Generative Genetic Algorithm Paradigm

**DOI:** 10.1101/2025.10.23.683781

**Authors:** Bliss Cui, Peter Bex

## Abstract

Central vision loss (CVL) from age-related macular degeneration (AMD) profoundly impacts daily life. While much research focuses on visual discrimination and reading, understanding how CVL affects emotional perception remains unexplored. We adapted the genetic-algorithm face paradigm introduced by Carlisi et al. (2021) and extended by Binetti et al. (2022), combining it with a three-dimensional morphable face model, to investigate how healthy young, healthy older, and CVL-affected individuals internally represent facial emotions. Participants evolved faces they perceived to best express 13 distinct emotions (or 4 for the older healthy group). Using a three-cohort design, we separated age effects from CVL-specific effects. Results revealed four significant group differences: Awe Convergence (H = 10.97, p = 0.001), Shame Intensity (H = 10.04, p = 0.005), Interest Range (H = 6.35, p = 0.016), and Interest Stability (H = 4.86, p = 0.026). Additionally, cosine similarity analysis in 199-dimensional face space showed cohorts use fundamentally different facial feature configurations (mean cosine ≈0.006 for YNG vs CVL), suggesting CVL may reorganize emotion representations rather than simply scaling them. These preliminary findings raise questions about whether all emotions require uniform amplification in CVL and point toward the potential value of selective rehabilitation over broad caricaturing approaches.

## Introduction

Age-related macular degeneration (AMD) is a leading cause of vision loss in older adults, affecting approximately 1 in 12 people over age 50 in developed countries (Wong et al., 2014). AMD typically causes central vision loss (CVL), impairing visual acuity and contrast sensitivity while leaving peripheral vision relatively intact. Most research on AMD has focused on visual tasks like reading, navigation, and face recognition (Logan et al., 2017; Boucart et al., 2008; Johnson et al., 2017; Wallis et al., 2014). However, emotion perception, the ability to judge emotional expressions on faces (Ekman & Friesen, 1971; Adolphs, 2002), is a critical social skill that may also be compromised in CVL, with potential cascading effects on social interaction and quality of life (Loughman & Flynn, 2009; Rovner & Casten, 2002; Lane et al., 2018).

To understand whether and how CVL affects emotion perception, we need methods that move beyond behavioral discrimination tasks. Internal representations, the mental models observers construct for different emotions, may reveal changes in how CVL affects social signal processing. Here we adopt the genetic-algorithm approach to internal emotion representation introduced by Carlisi et al. (2021) and extended by Binetti et al. (2022), in which participants iteratively evolve faces toward their internal representation of a target emotion. Building on our own prior work examining contextual and distortion effects on facial expression perception (Cui et al., 2023; Cui & Bex, 2022), we implement this approach within the 199-dimensional shape space of the Basel Face Model and apply it, for the first time, to individuals with central vision loss. This method captures the implicit, multidimensional structure of emotion categories (Barrett et al., 2011) rather than asking participants to explicitly rate stimuli.

A key challenge in studying age effects on emotion perception is that CVL is age-correlated: older AMD patients naturally have experienced longer exposure to central vision loss. Therefore, comparing only young and CVL groups confounds age with CVL status. We addressed this by recruiting three cohorts: young healthy (YNG, mean age 26–28 years), older healthy without AMD (OLD, mean age 72–75 years), and older CVL-affected (CVL, mean age 72–75 years with confirmed AMD). This three-cohort design allows us to statistically separate age effects from CVL-specific effects.

We hypothesized that CVL would alter the internal representations of emotions, with some emotions showing greater distortion than others. If visual information is differentially weighted in encoding different emotions (e.g., via distinct spatial frequency channels), we would expect selective rather than uniform changes across emotions.

## Methods

### Participants

Three cohorts were recruited: (1) YNG: 8 healthy young adults (mean age = 27.1 years, SD = 2.0; 3 male, 5 female); (2) OLD: 4 healthy older adults without AMD (mean age = 73.5 years, SD = 1.3; 1 male, 3 female); (3) CVL: 8 older adults with central vision loss from AMD (mean age = 73.9 years, SD = 7.0; 3 male, 5 female). All participants had corrected-to-normal vision (or best corrected with contact lenses) except for the CVL group, who had confirmed AMD diagnosed by an optometrist or ophthalmologist (Fletcher et al., 2012; Keane & Sadda, 2014). Inclusion criteria for the CVL group required best-corrected visual acuity of 20/60 or worse in at least one eye due to central scarring or drusenoid changes. Participants provided informed consent, and the study was approved by the Northeastern University Institutional Review Board.

### Stimulus Generation

Facial stimuli were generated using the Basel Face Model (BFM; Paysan et al., 2009), a three-dimensional morphable face model parameterized by principal components in shape and texture. BFM parameterizes facial shape morphology through principal components derived from 3D face scans. These parameters capture natural variation in facial structure (e.g., jaw width, cheekbone prominence, eye size). It is important to clarify that BFM does not directly manipulate gaze direction or head pose; all faces in this study were rendered at 0° rotation with a fixed forward-facing gaze. However, certain shape parameters affecting the eye and brow region could produce configurations that appear to imply different gaze directions or subtle orientation changes. Faces were rendered with neutral texture, displayed at approximately 10×10 cm on a monitor at 60 cm viewing distance, and presented in randomized order to prevent learning or expectancy effects.

### Genetic Algorithm Procedure

The genetic-algorithm procedure follows the approach described by Carlisi et al. (2021) and Binetti et al. (2022), implemented here with the Basel Face Model’s shape parameters rather than the 149-dimensional avatar expression space used in those studies. For each emotion, participants completed a 6-generation evolutionary task. In each generation, 12 face variants were displayed. Participants selected the face(s) they perceived to best express the target emotion. Selected faces served as parents for the next generation, with offspring created by interpolating parameters between parents and adding small random mutations (±5% of parameter range). This process iteratively evolved faces toward the participant’s internal representation of the emotion.

Each emotion trial (6 generations × 12 faces) took approximately 1.9–6.6 minutes to complete. Total session durations averaged 85.8 minutes (SD = 21.3) for YNG and 46.3 minutes (SD = 12.6) for CVL (both completing 13 emotions), and 7.5 minutes (SD = 1.5) for OLD (4 emotions). CVL participants completed trials faster per emotion (M = 3.6 min) than YNG (M = 6.6 min), possibly reflecting reduced deliberation time due to visual constraints or different selection strategies.

### Emotion Selection

YNG and CVL cohorts completed all 13 emotion trials: Amusement, Anger, Awe, Contempt, Disgust, Embarrassment, Fear, Happiness, Interest, Pride, Sadness, Shame, and Surprise. The OLD cohort completed four emotions (Amusement, Anger, Disgust, Happiness) based on their status as well-characterized basic emotions with established neural and behavioral signatures. This constraint was necessary due to fatigue and time considerations for older participants.

### Data Analysis

For each emotion, we extracted four metrics from the evolved faces: (1) Intensity: mean absolute deviation of evolved parameters from the population mean, quantifying how extreme the representation is; (2) Range: the spread of the evolved parameters (standard deviation); (3) Stability: coefficient of variation across the six generations; and (4) Convergence: the generation at which each participant reached 90% of maximum intensity. We performed Kruskal-Wallis (KW) tests to determine if any metric showed significant group differences (YNG vs. OLD vs. CVL). For comparisons with only two groups, we computed effect sizes (Cohen’s d) using pairwise means and pooled standard deviations. A face space analysis computed cosine similarity between cohort mean vectors in 199-dimensional BFM parameter space to determine whether cohorts represented emotions as scaled versions of a common template or as fundamentally different configurations.

## Results

### Convergence Analysis

Convergence analysis showed that most participants achieved 90% of maximum intensity well before generation 6. Across all participant-emotion combinations (n = 216), 73.1% (158/216) converged by generation 4, 84.3% (182/216) by generation 5, and 100% by generation 6. By cohort: YNG showed 71.2% convergence by gen 4, 83.7% by gen 5, and 100% by gen 6; OLD showed 50.0%, 87.5%, and 100%; CVL showed 76.9%, 84.6%, and 100%. These findings validate the 6-generation design as sufficient for the vast majority of cases, though adaptive stopping rules could improve efficiency in future studies.

### Expanded Evolutionary Metrics

To comprehensively characterize emotion evolution, we computed Range and Stability metrics for all emotion-cohort combinations. Range quantifies the spread (standard deviation) of evolved parameters; Stability measures consistency across generations. Kruskal-Wallis tests revealed that Range showed significant group differences only for Interest (H = 6.35, p = 0.016; all other emotions p > 0.05). Similarly, Stability showed significant differences only for Interest (H = 4.86, p = 0.026; all other emotions p > 0.05). This selective sensitivity demonstrates that Interest was the emotion most consistently affected by group membership, showing significant effects across Range, Stability, and large effect size differences in Intensity.

### Direction Analysis in Face Space

We computed cosine similarity between cohort mean vectors in the 199-dimensional face shape parameter space to determine whether groups construct emotions as simply more or less intense versions of a common template, or whether they use fundamentally different facial feature configurations. Results showed: YNG vs CVL (mean cosine = 0.006), YNG vs OLD (mean cosine = 0.017), and OLD vs CVL (mean cosine = −0.139). These near-zero to slightly negative values indicate that cohorts are approximately orthogonal (perpendicular) in face space, suggesting their emotion representations point in substantially different directions. Individual emotions ranged from −0.194 (Awe) to +0.205 (Fear) for YNG vs CVL comparisons. This finding suggests that CVL may not simply create more or less intense versions of the same emotion templates, but could involve different configurations of facial features.

### Key Group Differences

Four metrics reached statistical significance (Table 2): Awe Convergence (H = 10.97, p = 0.001), Shame Intensity (H = 10.04, p = 0.005), Interest Range (H = 6.35, p = 0.016), and Interest Stability (H = 4.86, p = 0.026). Table 3 presents pairwise effect sizes (Cohen’s d) for the significant comparisons.

**Table 1.** Participant Demographics.

| Cohort | N | Mean Age (SD) | Emotions Tested |
| --- | --- | --- | --- |
| YNG (Healthy Young) | 8 | 27.1 (2.0) | 13 |
| OLD (Healthy Older) | 4 | 73.5 (1.3) | 4 (basic) |
| CVL (AMD-Affected) | 8 | 73.9 (7.0) | 13 |

**Table 2.** Significant Kruskal-Wallis Tests (α = 0.05)

| Emotion | Metric | H-statistic | p-value |
| --- | --- | --- | --- |
| Awe | Convergence | 10.97 | 0.001 |
| Shame | Intensity | 10.04 | 0.005 |
| Interest | Range | 6.35 | 0.016 |
| Interest | Stability | 4.86 | 0.026 |

**Table 3.** Pairwise Effect Sizes (Cohen’s d) for Significant Comparisons.

| Emotion | YNG vs CVL | YNG vs OLD | OLD vs CVL | Magnitude |
| --- | --- | --- | --- | --- |
| Shame (Intensity) | −1.71 | N/A | N/A | Large |
| Anger (YNG vs OLD) | N/A | −0.96 | N/A | Large |
| Interest (Range) | −1.01 | N/A | N/A | Large |
| Awe (Convergence) | −0.42 | N/A | N/A | Small |

**Table 4.** Post Hoc Power Analysis Summary (Power = 0.80, α = 0.05)

| Comparison | Cohen's $d$ | $n$ per Group | Total $N$ |
| --- | --- | --- | --- |
| Shame (YNG vs CVL) | −1.71 | 6 | 12 |
| Anger (OLD vs CVL) | 1.31 | 10 | 20 |
| Interest (YNG vs CVL) | −1.01 | 16 | 32 |
| Anger (YNG vs OLD) | −0.96 | 18 | 36 |
| Amusement (YNG vs CVL) | 0.84 | 23 | 46 |

**Figure 1.**
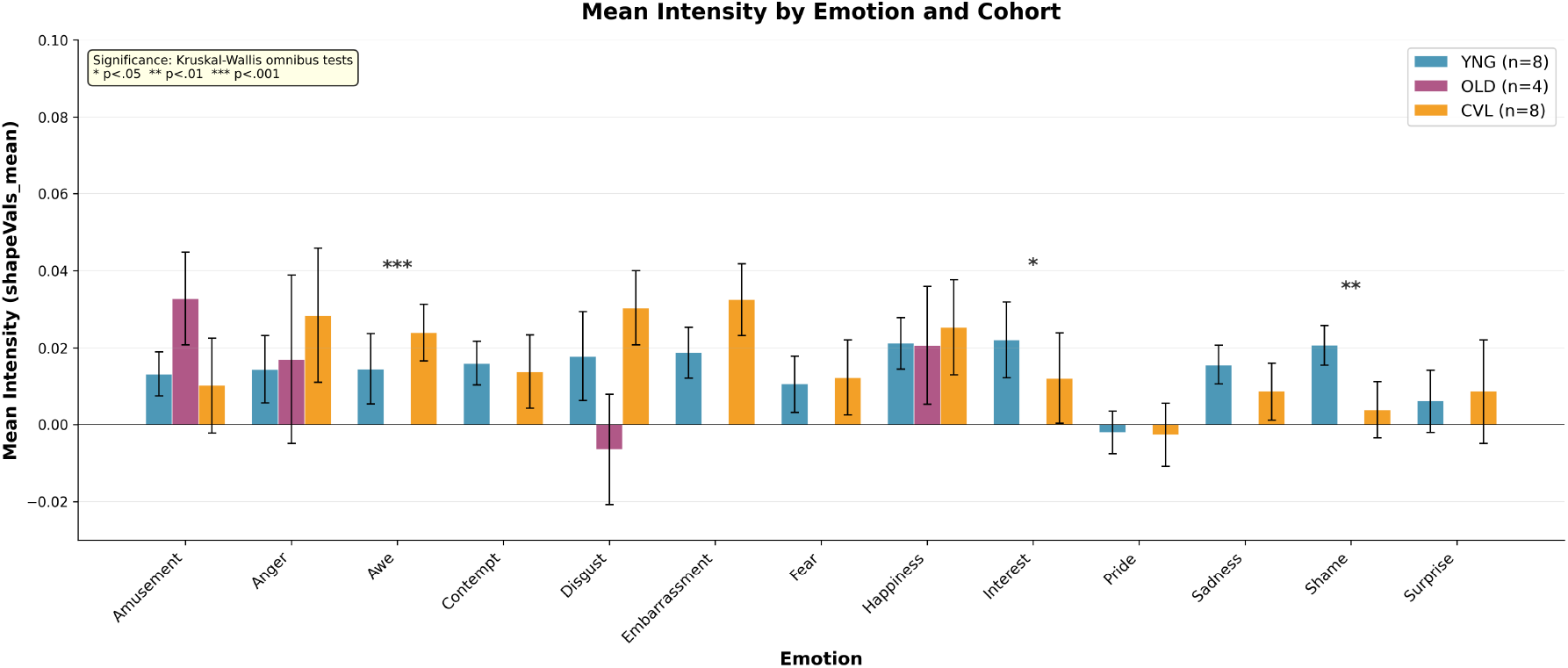
Mean Intensity by emotion and cohort. Error bars represent ±1 standard error. Asterisks indicate significant Kruskal-Wallis omnibus tests with permutation p-values (* p < .05, ** p < .01, *** p < .001). Note: Intensity metric shown here; Awe significance is for Convergence metric (not Intensity). Missing OLD bars indicate emotions not tested in that cohort.

**Figure 2.**
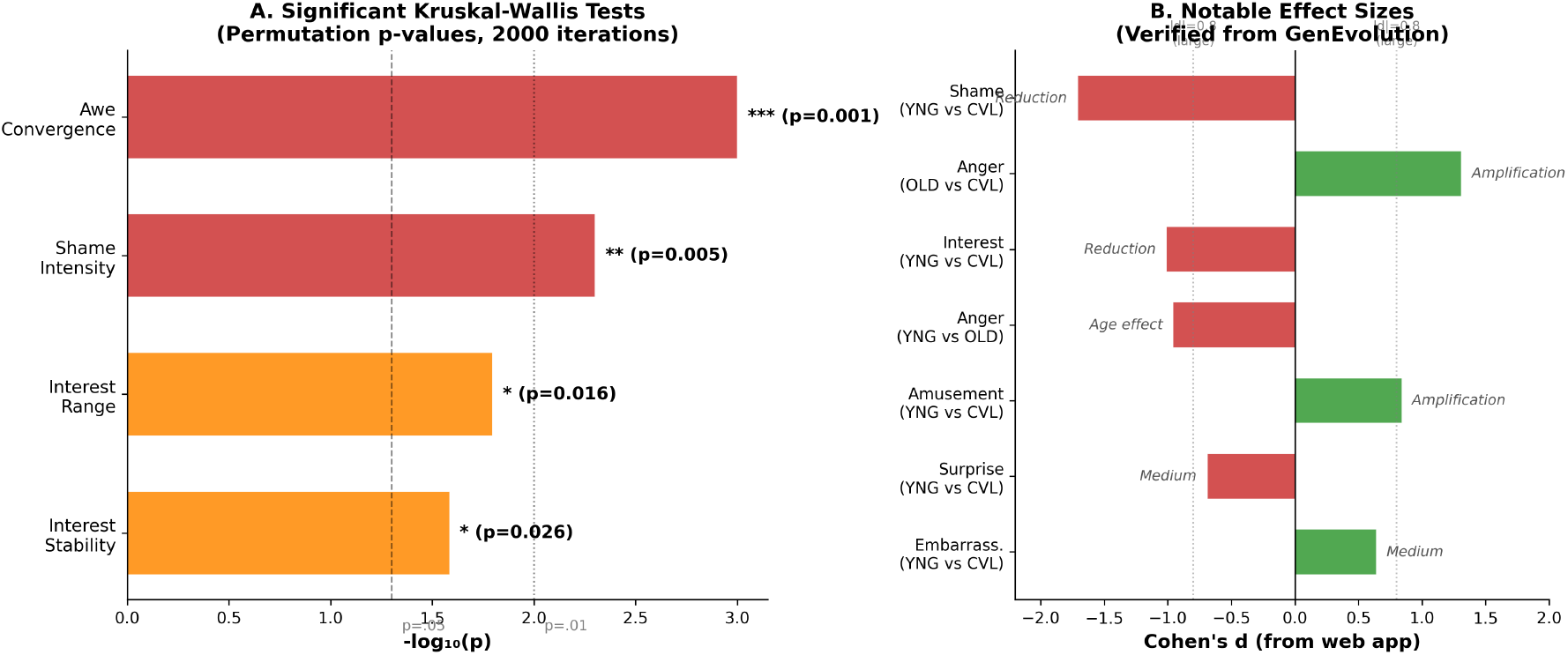
Summary of verified significant findings. (A) Kruskal-Wallis tests reaching significance, displayed as –log_10_(p). (B) Notable pairwise effect sizes, categorized by pattern type. Green bars = CVL higher (amplification); Red bars = YNG/OLD higher (reduction/age effect).

### Age-Related Patterns

For the four emotions tested in all three cohorts (Amusement, Anger, Disgust, Happiness), we observed mixed age patterns. Anger showed a large negative effect for YNG vs OLD (d = −0.96), with older adults showing higher intensity values. This pattern can be interpreted in two competing ways: (1) positivity bias hypothesis, in which older adults attenuate representations of negative emotions (consistent with socioemotional selectivity theory; Carstensen et al., 2006); or (2) heightened sensitivity hypothesis, in which older adults may require less facial exaggeration to internally perceive strong anger, reflecting a lower perceptual threshold or heightened sensitivity to negative affect. The current paradigm cannot distinguish between these interpretations, as the evolved faces reflect internal representations rather than explicit perceptual judgments. In contrast, Amusement, Disgust, and Happiness showed small effect sizes (|d| < 0.5), suggesting these emotions remain relatively preserved across age groups.

## Discussion

This study presents preliminary evidence that central vision loss may alter how individuals internally represent emotions. Using an evolutionary paradigm, we found significant group differences in four metrics (Awe Convergence, Shame Intensity, Interest Range, and Interest Stability), suggesting that CVL does not uniformly affect all emotions.

### Preservation of Most Emotion Representations

A striking finding is the preservation of most emotion representations in CVL. Seven of 13 emotions showed small or negligible effect sizes (|d| < 0.5) in YNG vs CVL comparisons: Contempt (−0.24), Disgust (+0.03), Fear (−0.10), Happiness (−0.12), Pride (−0.11), Sadness (−0.27), and Awe (−0.42). This suggests that CVL may leave most emotion representations largely intact. The alterations are concentrated in a subset of emotions: Shame (large reduction, d = −1.71), Interest (large reduction, d = −1.01), Surprise (medium reduction, d = −0.69), Amusement (amplified in CVL, d = +0.84 for YNG vs CVL), and Embarrassment (amplified, d = +0.64). This selective vulnerability is theoretically interesting: if emotions relied on a single common visual feature channel, we would expect uniform effects across all emotions. Instead, the differential impact suggests emotions recruit distinct combinations of visual features.

Clinically, this selective pattern has implications for rehabilitation. Rather than universal amplification (caricaturing) of all emotions, targeted interventions could focus on emotions most compromised in CVL (particularly Shame and Interest). In contrast, emotions that remain relatively preserved may not require intervention, potentially reducing cognitive load and improving compliance with vision rehabilitation programs.

### Reorganization Rather Than Simple Scaling

The face space direction analysis revealed near-zero cosine similarities between cohorts (mean ≈ for YNG vs CVL), indicating that emotion representations are not simply scaled up or down versions of a common template. Instead, CVL may reorganize which facial features are recruited for each emotion. If confirmed with larger samples, this pattern would suggest that CVL does not merely suppress or amplify visual signals, but may alter the multidimensional structure of emotion representation. One possible explanation is adaptive recalibration, in which CVL individuals learn to encode emotions using remaining visual channels or by relying more heavily on non-visual information (e.g., gaze direction implications from subtle shape changes).

### Convergence and Paradigm Efficiency

The convergence analysis demonstrates that 100% of participant-emotion combinations reached maximum intensity by generation 6, with 73% converging by generation 4. This supports the 6-generation design as likely sufficient for capturing the evolved representations. Future studies could implement adaptive stopping rules (automatically ending the trial when convergence is detected) to reduce session duration without sacrificing data quality. This would be particularly valuable for older and CVL participants, who may experience fatigue.

### Age Effects Revisited: Positivity Bias vs. Heightened Sensitivity

The large negative effect for Anger in YNG vs OLD (d = −0.96) can be interpreted in two competing frameworks. The positivity bias account suggests older adults actively suppress negative emotions in their internal models, consistent with socioemotional selectivity theory (Carstensen et al., 2006). However, an alternative interpretation (heightened sensitivity) proposes that older adults achieve the same perception of anger with less exaggeration, reflecting a lower threshold or increased sensitivity to negative affect. The genetic algorithm captures the internal representation that best expresses an emotion from the participant’s perspective, but it cannot distinguish whether reduced intensity reflects active suppression or heightened sensitivity. Behavioral discrimination tasks (e.g., asking whether a face is angry) would clarify this distinction in future work.

### Shame: CVL or Age Effect?

The largest effect size was for Shame Intensity (d = −1.71, YNG vs CVL), with CVL participants showing substantially reduced intensity. Shame involves gaze aversion and subtle facial cues (Keltner, 1995) that may be difficult to perceive with impaired central vision. However, we cannot definitively attribute this to CVL because Shame was not tested in the OLD cohort due to time constraints. Therefore, this effect could reflect age-related changes, CVL-specific changes, or their interaction. This is a key limitation and suggests that future studies should test all emotions across all cohorts when feasible.

### Implications for Face Caricaturing in AMD Rehabilitation

Recent work has shown that caricaturing (exaggerating facial features) improves facial expression recognition both in low-resolution images and in AMD patients (Lane et al., 2019). This success suggests that amplifying visual information can compensate for central vision loss. However, our preliminary findings raise questions about whether all emotions would benefit equally from uniform caricaturing. We found that some emotions in CVL are already amplified (Amusement: d = +0.84; Embarrassment: d = +0.64), while others are reduced (Shame: d = −1.71; Interest: d = −1.01). Uniform caricaturing could, in principle, further amplify emotions already exaggerated in CVL, while potentially helping emotions that are under-represented. A more targeted approach (selectively exaggerating emotions that are reduced while leaving preserved emotions unchanged) might warrant exploration in future rehabilitation research.

### Spatial Frequency Hypothesis and Future Directions

One theoretical explanation for the selective emotion effects is the spatial frequency hypothesis: different emotions may preferentially rely on different visual frequency bands (e.g., low spatial frequencies for broad facial shape; high frequencies for fine details like eye wrinkles; Liu et al., 2000; Vuilleumier et al., 2003). Since CVL preferentially affects central vision with high acuity, it may differentially impact emotions that rely on high-frequency information. To directly test this hypothesis, future research could compute the amplitude spectrum (via Fourier analysis) of evolved faces by emotion and cohort. Emotions showing larger shifts between YNG and CVL representations would exhibit stronger frequency-specific signatures.

### Eye Tracking and Gaze-Based Intervention

Understanding gaze patterns during emotion perception could reveal adaptive strategies. Recent work has shown that gaze-derived biofeedback, in which observers are shown explicit feedback based on their own attention patterns, can improve visual classification performance beyond training alone (Armengol-Urpi et al., 2026). If CVL individuals develop distinctive fixation strategies during emotion face evolution, these patterns could be quantified and potentially used to train healthy observers, or retrained in CVL individuals, to improve emotion perception efficiency. Future studies should include eye tracking to identify such strategies.

### Limitations and Power Analysis

The study has several important limitations. First, sample sizes were modest (YNG = 8, OLD = 4, CVL = 8), which limits statistical power for detecting smaller effects. A post hoc power analysis (assuming power = 0.80, α = 0.05, two-tailed) reveals the required sample sizes for our observed effect sizes: Shame (d = 1.71) required n = 6 per group; Anger OLD vs CVL (d = 1.31) required n = 10; Interest (d = 1.01) required n = 16; Anger YNG vs OLD (d = 0.96) required n = 18; Amusement (d = 0.84) required n = 23. Our sample sizes are adequate for the large effects we detected but would have limited power to detect medium or small effects. A follow-up study with 20–25 participants per cohort would provide robust detection of all large effects and enable more precise estimation of medium effects.

Second, the OLD cohort tested only four emotions due to time constraints, preventing us from isolating age effects for emotions like Shame and Interest. Third, while the genetic algorithm reveals internal representations, it does not directly measure behavioral emotion discrimination or provide insight into the cognitive strategies participants use. Fourth, the study is correlational; we cannot determine causal mechanisms (i.e., whether CVL produces the observed changes, or whether they reflect pre-existing differences or compensation strategies). Finally, generalizability is limited to individuals with AMD; results may not extend to other causes of central vision loss (e.g., Stargardt disease, cone-rod dystrophy).

## Conclusion

This study provides preliminary evidence that central vision loss from AMD may selectively alter internal emotion representations. Rather than uniform suppression or amplification, our results suggest that CVL could reorganize the facial features recruited for different emotions, as indicated by near-zero cosine similarities in high-dimensional face space. Most emotion representations appear preserved, but a subset (particularly Shame and Interest) show notable changes. These findings raise questions about universal caricaturing approaches and point toward the potential value of targeted, emotion-specific rehabilitation strategies, though confirmation with larger samples is needed. Future research should employ larger samples, test all emotions across all cohorts, incorporate behavioral discrimination tasks, and use eye tracking to identify gaze-based intervention targets.

## Acknowledgements

The authors thank Nicole Ross and Olivia Wynn for their contributions to participant recruitment and data collection.

## AI Statement

An AI assistant (Claude, Anthropic) was used to assist with manuscript preparation, including formatting, proofreading, and statistical verification against pre-existing analysis outputs. All scientific content, study design, data collection, analysis, and interpretation were performed by the authors.

## Data Availability

The data and analysis code supporting this study are available on GitHub upon reasonable request to the corresponding author.

## Author Contributions

Bliss Cui: Conceptualization, Methodology, Software, Formal Analysis, Investigation, Data Curation, Writing – Original Draft, Writing – Review and Editing, Visualization, Project Administration. Peter Bex: Conceptualization, Methodology, Supervision, Resources, Writing – Review and Editing, Funding Acquisition.

## Ethical Approval

This study was approved by the Northeastern University Institutional Review Board. All procedures were conducted in accordance with the Declaration of Helsinki.

## Consent to Participate

All participants provided written informed consent prior to participation in this study.

